# Functional Deorphanization and Subtype-Selective Pharmacology of Three Tyramine Receptors in the Disease Vector, *Aedes aegypti*

**DOI:** 10.64898/2026.07.24.740580

**Authors:** Salwa Afifi, Jean-Paul Paluzzi

## Abstract

Biogenic amines such as tyramine (TA) and octopamine (OA) are central regulators of insect physiology and behaviour, acting through G protein–coupled receptors (GPCRs) to control reproduction, locomotion, metabolism, olfaction and hydromineral homeostasis. Although TA was once considered solely a biosynthetic precursor to OA, it is now recognized as an independent signaling molecule acting through distinct tyramine receptors (TARs). Owing to their invertebrate-specific roles and absence in vertebrates, TARs represent promising molecular targets for selective insecticide development. In the mosquito *Aedes aegypti*, a major arboviral vector of dengue and Zika viruses, the functional and pharmacological properties of TARs have not been characterized. Here, we functionally deorphanized and comparatively characterized three putative *A. aegypti* tyramine receptors (AaTAR1–AaTAR3) using a heterologous assay, revealing subtype-specific pharmacological profiles and antagonist sensitivities. All three receptors were robustly activated by TA in a concentration-dependent manner, whereas OA exhibited significantly lower potency on each receptor subtype, consistent with a strong preference for TA. All three TARs were unresponsive to dopamine and serotonin, even when using supraphysiological concentrations, indicating high ligand specificity. Antagonist profiling revealed pronounced subtype-specific pharmacology: yohimbine strongly suppressed AaTAR1, phentolamine most effectively inhibited AaTAR2, whereas AaTAR3 exhibited reduced sensitivity to several classical aminergic antagonists, suggesting a pharmacologically distinct subtype. These findings establish that AaTAR1, AaTAR2 and AaTAR3 are bona fide functional tyramine receptors, define their ligand selectivity and subtype-specific pharmacology, and provide a comparative framework for understanding mosquito tyraminergic signaling, highlighting their potential as targets for next-generation vector control strategies.

## 1. Introduction

Despite decades of intensive research and control efforts, mosquito-borne arboviral infections remain a major and growing hazard to worldwide public health. The mosquito *Aedes aegypti* is a known vector of dengue, Zika, chikungunya and yellow fever viruses and is highly adapted to urban environments, enabling close association with human populations (Kotsakiozi et al., 2017; Leta et al., 2018; McGregor and Connelly, 2021; Neiderud, 2015; Patterson et al., 2016; Powell and Tabachnick, 2013; Saiz et al., 2016). Dengue virus alone is estimated to infect almost 390 million people annually, with nearly 96 million clinically apparent cases, therefore affecting nearly half of the world’s population at risk of infection (Bhatt et al., 2021, 2013; Hasan et al., 2016; Jing and Wang, 2019; Otu et al., 2019). In addition, Zika virus infection during pregnancy is associated with congenital Zika syndrome, including severe neurological defects such as microcephaly (Huang et al., 2016; Krauer et al., 2017). The distribution of *A. aegypti* worldwide has been driven by urbanization, global climate change and increased human travel, worsening the burden of mosquito-borne diseases (Burt et al., 2017; Liu-Helmersson et al., 2019; Sakkas et al., 2016). Although current control strategies (e.g., insecticides, vaccines, genetic approaches and *Wolbachia*-based interventions) have contributed towards new disease management strategies, their long-term effectiveness is limited by the rapid emergence of insecticide resistance and operational constraints (McGregor and Connelly, 2021; Weeratunga et al., 2017). All of these challenges require urgency in identifying novel molecular targets for innovative vector control approaches.

Biogenic amines, which include tyramine (TA), octopamine (OA), serotonin and dopamine, function as key neurotransmitters, neuromodulators and neurohormones in insects, regulating a variety of physiological processes such as reproduction, metabolism, locomotion as well as ionic and osmotic homeostasis, mainly by the activation of G protein–coupled receptors (GPCRs) (Evans, 1980; Roeder, 2005; Verlinden et al., 2010). In invertebrates, TA and OA act as important monoaminergic signaling molecules with functions analogous to adrenaline and noradrenaline in vertebrates (Fuchs et al., 2014; Roeder, 2020, 2005, 1999). Both TA and OA are invertebrate-specific signaling systems; they represent attractive targets for selective insecticides (Blenau and Baumann, 2001; Jonsson and Hope, 2007). Indeed, TA and OA receptors are established targets of insecticidal and acaricidal compounds, including the formamidine amitraz, which primarily targets OA receptors but has also been shown to act on a subset of tyramine receptors (Jonsson and Hope, 2007; Kumar, 2019; Wu et al., 2013). Furthermore, natural products (e.g., monoterpenes) have been reported to disrupt OA and TA signaling in several insects, which could thereby enhance the translational potential of these aminergic pathways for vector control (Finetti et al., 2020, 2021b; Jankowska et al., 2018).

TA is synthesized from tyrosine by tyrosine decarboxylase and can be hydroxylated to OA via tyramine β□hydroxylase (Lange, 2009; Roeder et al., 2003). While TA was earlier regarded simply as a biosynthetic precursor of OA, it is now established as a signaling entity that operates through specialized tyramine receptors (TARs) to mediate many physiological functions (Lange, 2009; Roeder, 2020). TARs have been identified across multiple insect species and are classified into three major subtypes (TAR1, TAR2 and TAR3), and each receptor displays its distinct pharmacological and functional properties. TAR1 has been extensively characterized and displays high affinity for TA, whereas TAR2 and TAR3 remain comparatively less well understood and may exhibit subtype-specific signaling and ligand sensitivity (Blenau and Baumann, 2001; Finetti et al., 2021a; Hana and Lange, 2017a; Ohta and Ozoe, 2014; Reim et al., 2017; Wu et al., 2015). Recent analyses using RT-qPCR have identified three putative TARs in *A. aegypti* with differential expression across developmental stages and tissues, including reported enrichment of TAR2 in ovaries and TAR3 in Malpighian tubules, suggesting their potential roles in reproduction and osmoregulation, respectively (Finetti et al., 2023). However, this study was limited to molecular characterization through expression profiling, while the ligand specificity, signaling properties and pharmacological profiles of these receptors have not been functionally characterized. Although other biogenic amines such as serotonin and dopamine have been implicated in regulating blood feeding behaviour and development in *A. aegypti* (Andersen et al., 2006; Ngai et al., 2019; Vinauger et al., 2018), the physiological roles and receptor-level pharmacology of TA and OA signaling in this species remain poorly understood. Here, we report the functional deorphanization and pharmacological characterization of three tyramine receptor subtypes (AaTAR1, AaTAR2 and AaTAR3) from *A. aegypti*. Ligand selectivity across biogenic amines and subtype-specific pharmacological profiles was defined using a panel of antagonists by heterologous expression in mammalian cells, coupled with an aequorin-based calcium mobilization assay. Our results provide the first comprehensive functional characterization of three TARs in *A. aegypti*, revealing distinct pharmacological signatures and providing a framework for the potential future development of subtype-selective modulators targeting aminergic signaling in mosquito vectors.

## 2. Materials and Methods

### 2.1. Experimental Animals and Rearing Conditions

A laboratory colony of *Aedes aegypti* (Liverpool strain) was maintained in the Department of Biology, York University (Toronto, ON, Canada). Semi-desiccated eggs were hatched in double-distilled water, and larvae were reared on a diet of 2% (w/v) beef liver powder and 2% (w/v) brewer’s yeast dissolved in water. Pupae were transferred to distilled water for adult emergence. All post-embryonic life stages were maintained at 26 °C under a 12:12 h light:dark photoperiod. Adult male and female mosquitoes were provided with continuous access to 10% (w/v) sucrose solution *ad libitum* via cotton wicks. For colony maintenance, adult females were offered the opportunity to blood-feed every 48 h using sheep’s blood in Alsever’s solution (Cedarlane Laboratories Ltd., Burlington, ON, Canada) delivered via an artificial membrane feeding system as described previously (Afifi et al., 2023; Rocco et al., 2017; Wahedi and Paluzzi, 2018).

### 2.2. Bioinformatic Predictions of the Topology and Post-Translational Modification Sites of Three TARs

The deduced amino acid sequences of AaTAR1, AaTAR2 and AaTAR3 were used for in silico prediction of receptor topology and putative post-translational modification sites. Transmembrane topology was predicted using the Constrained Consensus TOPology prediction server (CCTOP; https://cctop.ttk.hu/) (Dobson et al., 2015), and the final CCTOP consensus topology was used to define the predicted transmembrane domains, extracellular regions and intracellular regions. Potential N-linked glycosylation sites were predicted using NetNGlyc 1.0 (https://services.healthtech.dtu.dk/services/NetNGlyc-1.0/) (Gupta and Brunak, 2002). Only positive NetNGlyc-predicted Asn-Xaa-Ser/Thr motifs located within CCTOP-predicted extracellular regions were mapped onto the receptor topology. Potential phosphorylation sites were predicted using NetPhos 3.1 for serine, threonine and tyrosine residues (https://services.healthtech.dtu.dk/services/NetPhos-3.1/) (Blom et al., 2004, 1999). For figure visualization, only NetPhos-predicted intracellular phosphorylation sites with prediction scores ≥ 0.75 were displayed. The predicted topology and putative post-translational modification sites were visualized using Protter (https://protter.ethz.ch/#) (Omasits et al., 2014).

### 2.3. RNA Isolation, Primer Design, cDNA Synthesis and Cloning

Three putative tyramine receptor genes were selected for cloning from adult *A. aegypti*: AaTAR1 (XM_001652205.3; AAEL006844), AaTAR2 (XM_021837305.1; AAEL004396) and AaTAR3 (XM_021837306.1; AAEL004398). Total RNA was extracted from 1–4-day-old adult *A. aegypti* using the Monarch Total RNA Miniprep Kit (New England Biolabs, Whitby, ON, Canada) following the manufacturer’s instructions. RNA concentration and purity were assessed by micro-volume spectrophotometry using a Synergy 2 Multi-Mode Microplate Reader equipped with a Take3 micro-volume plate (BioTek, Winooski, VT, USA). For the initial cDNA amplification and sequence verification, gene-specific primers for AaTAR1, AaTAR2 and AaTAR3 were designed de novo using Primer3Plus implemented in Geneious Bioinformatics Software (Biomatters Ltd., Auckland, New Zealand). Primers were positioned in the 5′ and 3′ untranslated regions (UTRs) outside the predicted open reading frame (ORF) flanking the entire coding sequence (CDS) (**Table 1**).

**Table 1.**
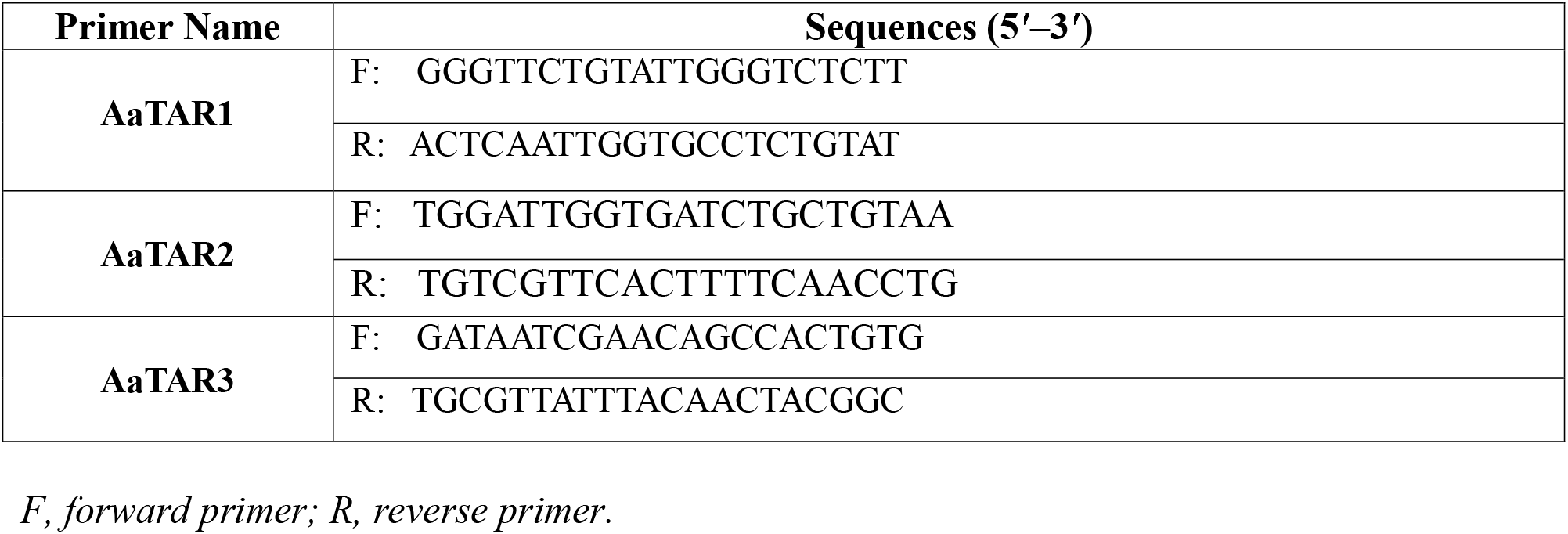
Primers used for initial amplification and cloning of three *A. aegypti* tyramine receptors into the pCE2 TA/Blunt-Zero vector.

First-strand cDNA was synthesized from ∼500 ng total RNA using SuperScript IV reverse transcriptase (Invitrogen, Thermo Fisher Scientific, ON, Canada) following the manufacturer’s guidelines, with gene-specific reverse primers for AaTAR1, AaTAR2 and AaTAR3 at a final concentration of 2 µM. The resulting cDNA was used as a template for PCR amplification of AaTAR1–AaTAR3 using Q5® High-Fidelity DNA Polymerase (New England Biolabs) with the target-specific forward and reverse primers listed in **Table 1**. PCR reactions were performed under the following cycling conditions: initial denaturation at 98 °C for 30 s, followed by 40 cycles of denaturation at 98 °C for 10 s, annealing at 64 °C for 20 s for all three receptors and extension at 72 °C for 1 min, with a final extension at 72 °C for 2 min. PCR products of the expected size were gel-purified and cloned into the pCE2 TA/Blunt-Zero vector using the 5-minute TA/Blunt-Zero Cloning Kit (GeneBioSystems, Burlington, ON, Canada). Plasmid DNA was purified using the Monarch® Plasmid Miniprep Kit (New England Biolabs, Whitby, ON, Canada), and insert sequences were verified by Sanger sequencing (Centre for Applied Genomics, Hospital for Sick Children, Toronto, ON, Canada) and whole-plasmid sequencing performed by Plasmidsaurus (South San Francisco, CA USA) using Oxford Nanopore Technology with custom analysis and annotation.

### 2.4. Preparation of Mammalian Expression Constructs for AaTAR1–AaTAR3

For functional heterologous expression in mammalian cells, a second set of primers was designed to amplify the verified full-length ORFs of AaTAR1, AaTAR2 and AaTAR3 for directional cloning into the pcDNA3.1^+^ expression vector (Life Technologies, Burlington, ON, Canada). These primers incorporated a Kozak consensus sequence (GCCACC) immediately upstream of the start codon, together with 5′ HindIII and 3′ XbaI restriction sites flanking each receptor ORF (**Table 2**). Full-length ORFs were amplified using Q5® High-Fidelity DNA Polymerase (New England Biolabs, Whitby, ON, Canada) with annealing temperatures of 66 °C for AaTAR1, 68 °C for AaTAR2 and 67 °C for AaTAR3. The amplified products and pcDNA3.1^+^ were restriction-digested, purified and ligated to generate mammalian expression constructs for downstream heterologous expression and pharmacological characterization. Construct orientations and sequence fidelity were confirmed by Sanger sequencing (Centre for Applied Genomics, Hospital for Sick Children, Toronto, ON, Canada) and whole-plasmid sequencing performed by Plasmidsaurus (South San Francisco, CA USA) using Oxford Nanopore Technology with custom analysis and annotation. Endotoxin-free, transfection-grade plasmid DNA was prepared from overnight bacterial cultures using the ZymoPURE™ II Plasmid Midiprep Kit (Zymo Research, Irvine, CA, USA) and used for subsequent mammalian cell transfection assays.

**Table 2.**
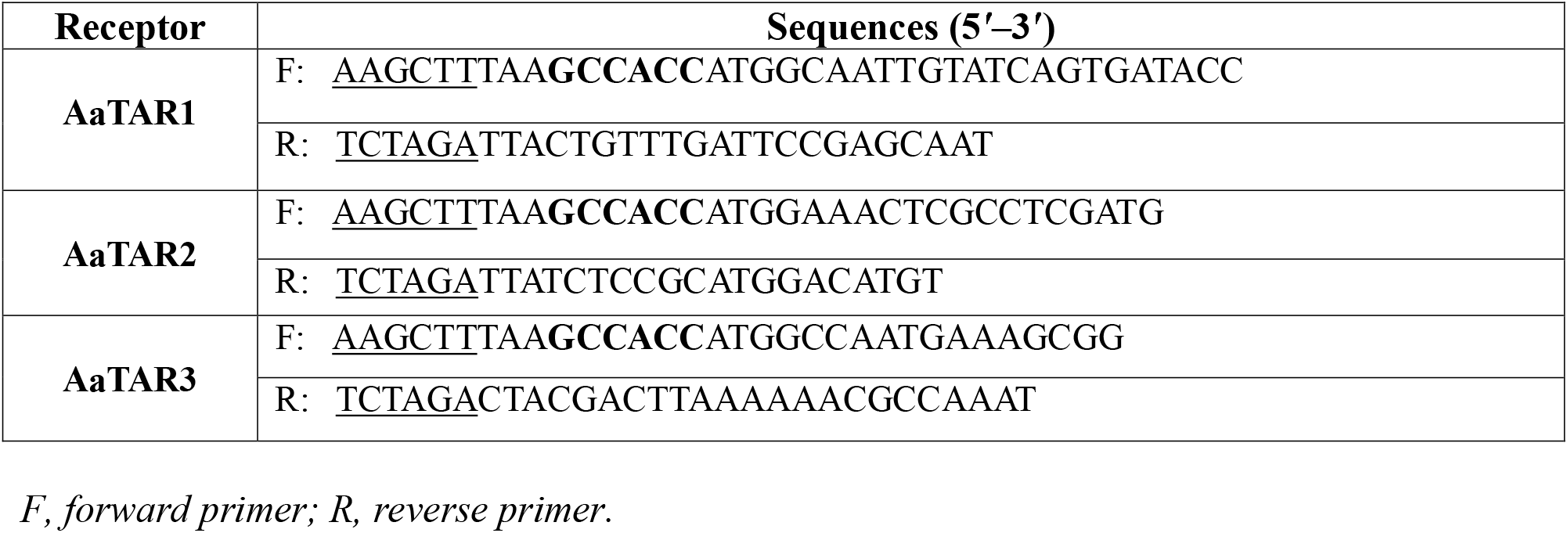
Primers used for directional subcloning *A. aegypti* tyramine receptor open reading frames (ORFs) (AaTAR1–3) into pcDNA3.1^+^ for mammalian expression. Kozak sequences are shown in bold; restriction sites (HindIII = AAGCTT; XbaI = TCTAGA) are underlined; F, forward primer; R, reverse primer.

### 2.5. Heterologous Functional Receptor Assays

#### 2.5.1 Chemicals

All biogenic amines (tyramine hydrochloride, (±)-octopamine hydrochloride, dopamine hydrochloride and serotonin hydrochloride) were prepared as 0.5 M stock solutions in double-distilled water. Biogenic amine antagonists (yohimbine hydrochloride, phentolamine hydrochloride, metoclopramide hydrochloride, gramine, epinastine hydrochloride and mianserin hydrochloride) were prepared in double-distilled water or DMSO as 25–50 mM stock solutions. Working solutions were freshly prepared in assay medium, and the final solvent concentration in experimental treatments was maintained at ≤0.1%. All chemicals were purchased from Sigma-Aldrich (Oakville, ON, Canada).

#### 2.5.2 Cell Culture and Transfection

A recombinant Chinese hamster ovary (CHO-K1) cell line stably expressing the bioluminescent calcium sensor aequorin (Gondalia et al., 2016; Paluzzi et al., 2012) was maintained in DMEM/F12 basal medium (Wisent Inc., St. Bruno, QC, Canada) supplemented with 10% heat-inactivated fetal bovine serum, 250 µg/mL Geneticin and 1× Antibiotic–Antimycotic (Thermo Fisher Scientific, Mississauga, ON, Canada) as reported previously (Oryan et al., 2018; Wahedi and Paluzzi, 2018). Cells were cultured at 37 °C in 5% CO□and transfected at ∼90–95% confluency with pcDNA3.1^+^ constructs encoding AaTAR1–AaTAR3 using PolyJet™ *in vitro* DNA transfection reagent (FroggaBio Inc., Concord, ON, Canada). Functional assays were performed ∼48 h post-transfection.

#### 2.5.3 Calcium Mobilization Assay

Transfected cells were harvested with DPBS-EDTA, resuspended at 1 × 10□cells/mL in BSA medium (DMEM:F12, 0.1% BSA, 1× Antibiotic-Antimycotic) and then incubated with 5 µM coelenterazine-h (NanoLight Technology, Pinetop, AZ, USA) for 3–4 h in the dark, following established protocols (Oryan et al., 2018). Cell suspensions were then diluted 10-fold in BSA medium and incubated for an additional 45-60 min. Assays were performed in white 96-well plates (Greiner Bio-One, Monroe, NC, USA). Luminescence was recorded for 20 s using a Synergy 2 Multi-Mode Microplate Reader equipped with an automated injector (BioTek, Winooski, VT, USA), following injection of 50 µL of cell suspension per well.

For the dose–response experiments, serial dilutions (10^-3^ to 10^-12^ M) of tyramine or octopamine were applied. Luminescence values were background-subtracted using BSA-only controls and normalized to the maximal ATP response (100 µM) within each independent experiment.

For agonist selectivity profiling, cells expressing AaTAR1–AaTAR3 were stimulated with tyramine, octopamine, dopamine or serotonin (10^-5^ M). In co-application experiments, either octopamine, dopamine or serotonin (10^-5^ M) was applied in combination with tyramine (10^-5^ M). For pharmacological characterization, tyramine (10^-5^ M) was pre-mixed with individual antagonists (10^-5^ M) in assay medium prior to cell injection. Receptor-expressing cells were then injected into individual wells containing the ligand/antagonist mixture, and luminescence was recorded as described above. Responses were normalized to the tyramine-only treatment (10^-5^ M), defined as 100% for each receptor subtype within each independent experiment. Negative controls consisted of wells containing BSA assay medium without agonist stimulation, whereas 100 µM ATP was used as a positive control to normalize the bioluminescent response. In addition, non-transfected CHO-K1 cells were tested with TA, OA and other biogenic amines to confirm that the compounds themselves did not induce a response in the absence of AaTAR expression. All experiments were performed in technical triplicate across at least three independent biological replicates.

### 2.6 Data Analysis

All data were analyzed using GraphPad Prism 9.0 (GraphPad Software, San Diego, USA). Dose– response curves and EC_50_ values were obtained by nonlinear regression using a sigmoidal dose– response model. Biogenic amine activation, co-application experiments and antagonist profiling were analyzed using one-way ANOVA followed by Tukey’s multiple comparisons test. Differences were considered statistically significant at p < 0.05.

## 3. Results

### 3.1 Cloning, Sequence Verification and Predicted Topology of Three *A. aegypti* Tyramine Receptors

Sequence verification confirmed the coding sequences of AaTAR1, AaTAR2 and AaTAR3, which were deposited in GenBank under accession numbers AaTAR1 (PZ585254), AaTAR2 (PZ585255) and AaTAR3 (PZ585256), respectively.

Bioinformatic analysis of the deduced amino acid sequences predicted that all three AaTARs possess the canonical seven-transmembrane architecture characteristic of G protein-coupled receptors, with an extracellular N-terminus and an intracellular C-terminus (**Fig. 1**). The deduced proteins consisted of 522 amino acids for AaTAR1 (**Fig. 1A)**, 720 amino acids for AaTAR2 (**Fig. 1B)** and 543 amino acids for AaTAR3 (**Fig. 1C)**. With respect to the putative post-translational modification sites, NetNGlyc predicted four extracellular N-linked glycosylation sites in each receptor. NetPhos predicted multiple intracellular phosphorylation sites within intracellular loops and the C-terminal cytoplasmic regions, with 15 sites in AaTAR1, 23 sites in AaTAR2 and 22 sites in AaTAR3. Together, these results indicate that AaTAR1, AaTAR2 and AaTAR3 encode GPCR-like receptors with conserved seven-transmembrane topologies and predicted extracellular glycosylation and intracellular phosphorylation sites.

**Figure 1.**
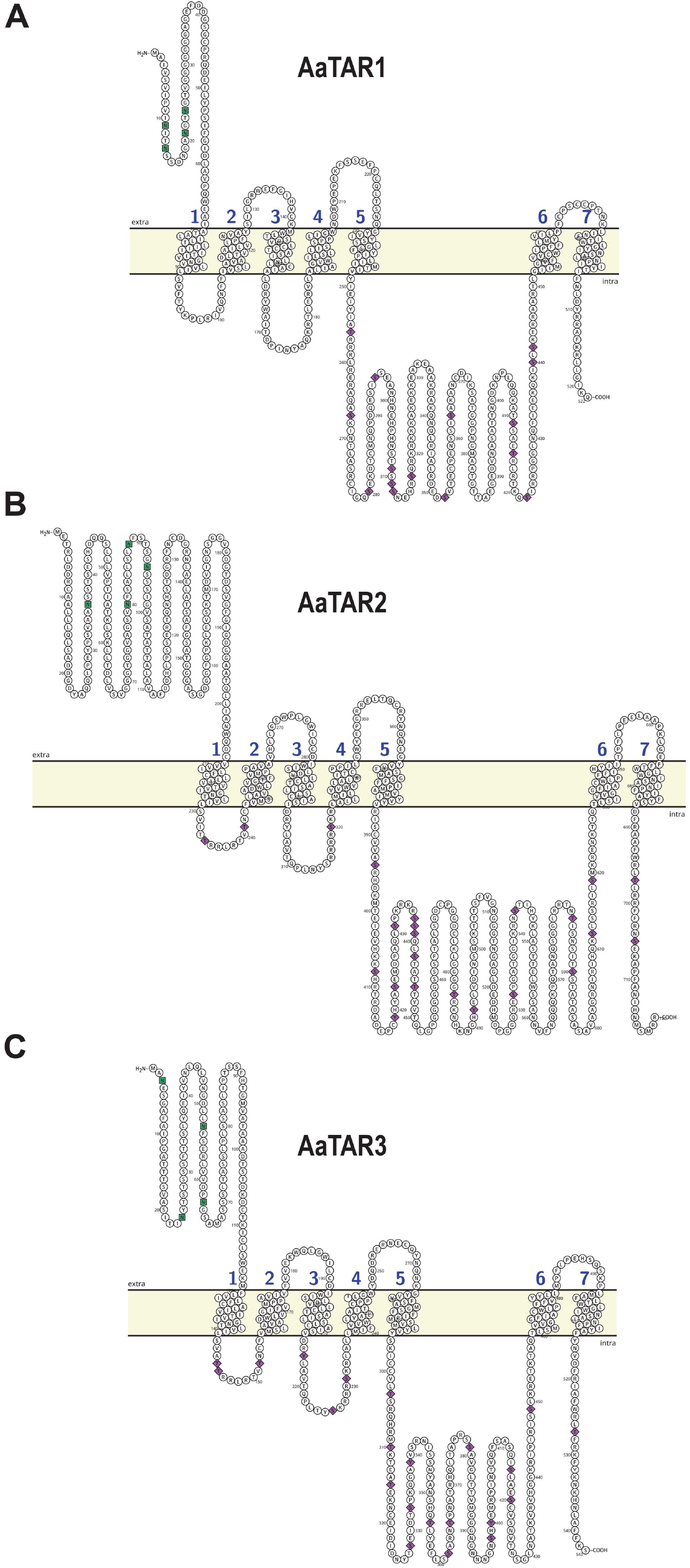
Predicted membrane topology and post-translational modification sites of three *Aedes aegypti* tyramine receptors AaTAR1, AaTAR2 and AaTAR3. The deduced amino acid sequences of AaTAR1 **(A)**, AaTAR2 **(B)** and AaTAR3 **(C)** were analyzed using CCTOP to predict transmembrane topology and receptor orientation. All three receptors were predicted to contain seven-transmembrane domains, with extracellular N-termini and intracellular C-termini. Potential N-linked glycosylation sites located in extracellular regions are indicated by green squares. Predicted phosphorylation sites located in intracellular regions are indicated by purple diamonds.

### 3.2 Agonist Selectivity Profiling of AaTAR1–AaTAR3

To evaluate ligand selectivity, AaTAR1, AaTAR2 and AaTAR3 were transiently expressed in CHO-K1 cells stably expressing aequorin (Gondalia et al., 2016; Paluzzi et al., 2012) and stimulated with the biogenic amines tyramine (TA), octopamine (OA), dopamine and serotonin (10^-5^ M). Luminescent responses were normalized to the TA (10^-5^ M) response within each receptor subtype. TA elicited the highest response across all three receptor subtypes, whereas OA produced marginal, receptor-dependent responses, albeit significantly higher compared to control receiving BSA media alone (**Fig. 2A–C**). Dopamine and serotonin did not elicit responses by any of the three receptors greater than the background luminescence observed in controls receiving BSA media alone. Overall, these results indicate a clear ligand preference for TA across all three AaTAR subtypes (**Fig. 2**).

**Figure 2.**
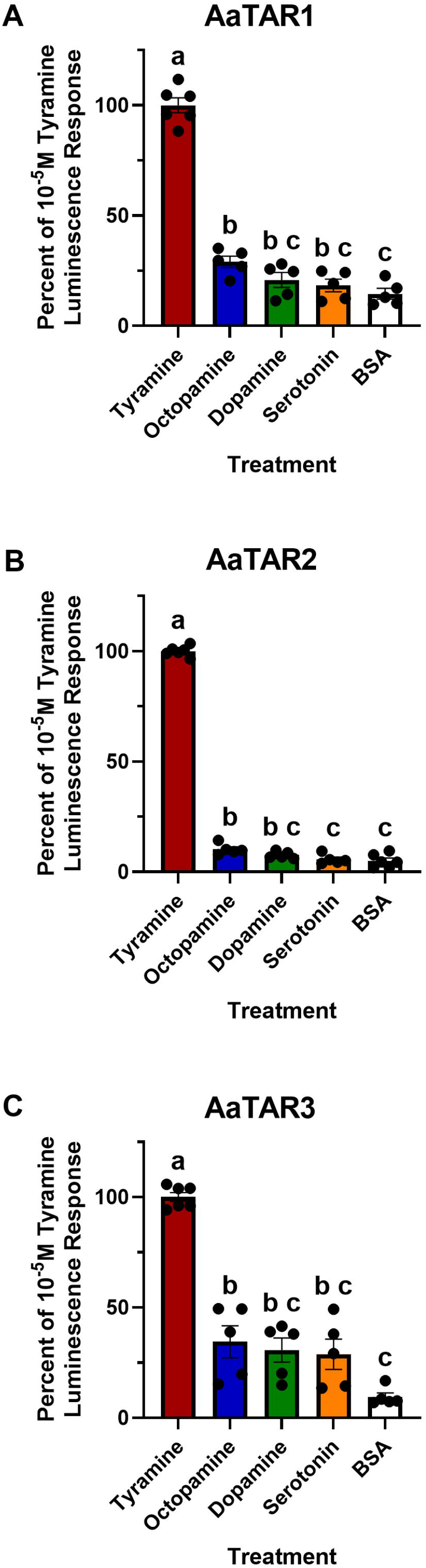
Biogenic amine–induced activation of AaTAR1–AaTAR3. Aequorin-CHO-K1 cells transiently expressing **(A)** AaTAR1, **(B)** AaTAR2 or **(C)** AaTAR3 were treated with tyramine (TA), octopamine, dopamine or serotonin (each 10^-5^ M). Luminescence responses were normalized to the TA response (10^-5^ M) within each receptor subtype (TA = 100%); BSA (vehicle) served as the negative control. TA produced the highest luminescence response across all receptor subtypes, whereas the other biogenic amines elicited lower, subtype-dependent responses. Different letters above bars indicate statistically significant differences among treatments within each receptor subtype as determined by one-way ANOVA followed by Tukey’s multiple-comparisons test; p < 0.05. Data represent mean ± SEM (n = 5–6).

### 3.3. Dose-Dependent Activation and Potency Profiling of AaTAR1–AaTAR3

To quantitatively assess receptor sensitivity and ligand potency, concentration–response analyses were performed for TA and OA across AaTAR1, AaTAR2 and AaTAR3 (**Fig. 3**). All three receptors exhibited sigmoidal, concentration-dependent activation in response to both ligands. TA produced more robust activation and exhibited greater potency than OA across all three receptor subtypes (**Fig. 3A–C**). In contrast, OA required substantially higher concentrations to achieve comparable responses. Nonlinear regression analysis revealed consistently lower EC_50_ values for TA than for OA across all three TAR subtypes (**Table 3**). Across receptor subtypes, TA was 271–496-fold more potent than OA (**Table 3**), confirming its greater ligand potency. To compare TA-evoked response magnitude, fold change in calcium–dependent luminescence at 10^-3^ M TA was evaluated across AaTAR1–AaTAR3 (**Fig. 3D**). AaTAR2 produced a significantly greater fold change in calcium-dependent luminescence than AaTAR1 and AaTAR3.

**Figure 3.**
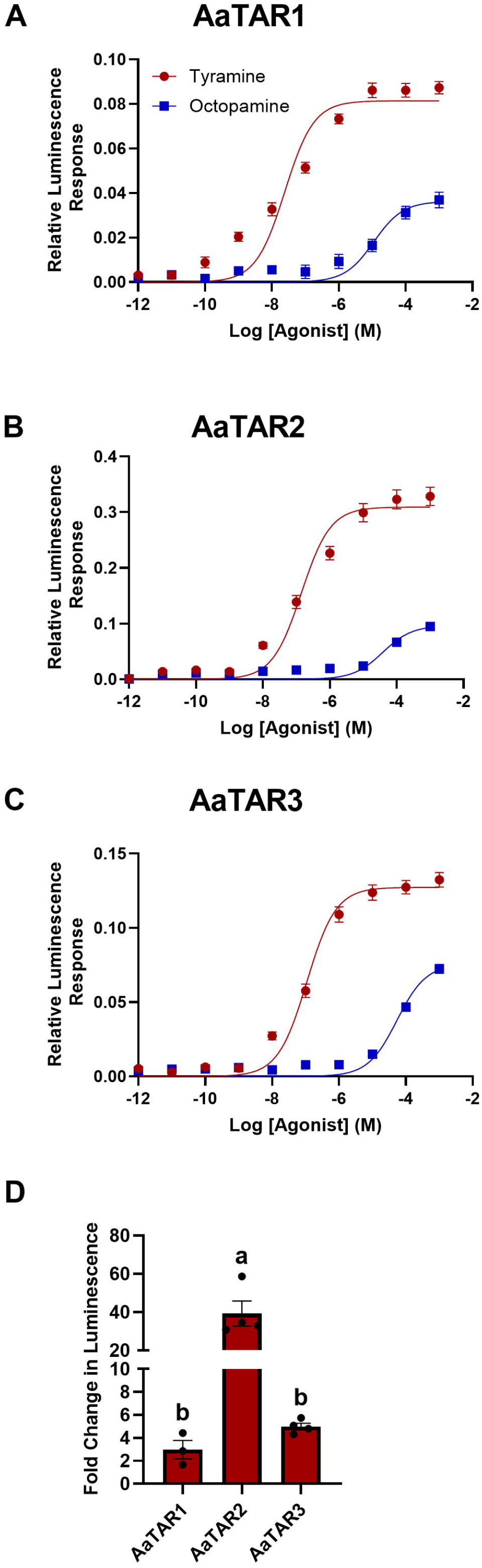
Dose–response analysis of tyramine and octopamine at AaTAR1–AaTAR3 and comparison of TA-evoked calcium response magnitude. Concentration–response curves were generated in CHO-K1 cells stably expressing aequorin and transiently expressing (**A**) AaTAR1, (**B**) AaTAR2 or (**C**) AaTAR3. Cells were stimulated with tyramine (TA; red circles) or octopamine (OA; blue squares) across a concentration range of 10^-12^–10^-3^ M. Luminescence responses are plotted as relative luminescence versus log□□agonist concentration (M), background-subtracted and normalized to the maximal ATP response (100 µM) within each independent experiment. TA produced higher maximal responses and had greater potency than OA on all three receptor subtypes. (**D**) Comparison of TA-evoked fold change in calcium-dependent luminescence across AaTAR1–AaTAR3 at 10^-3^ M TA. Fold change was calculated relative to the mean BSA background response. AaTAR2 showed a significantly greater fold change in calcium-dependent luminescence than AaTAR1 and AaTAR3. For panel D, differences among receptor subtypes were analyzed using one-way ANOVA followed by Tukey’s multiple-comparisons test. Different letters indicate significant differences among receptor subtypes (p < 0.05). Data represent mean ± SEM (n = 3–4).

**Table 3.** EC_50_ values and relative potency of octopamine versus tyramine at AaTAR1– AaTAR3.

| Receptor | Tyramine EC<sub>50</sub> (M)* | Octopamine EC<sub>50</sub> (M)* | Fold Difference (OA/TA) |
| --- | --- | --- | --- |
| **AaTAR1** | 2.447×10<sup>-8</sup> | 1.125×10<sup>-5</sup> | 460 |
| **AaTAR2** | 1.414×10<sup>-7</sup> | 3.834×10<sup>-5</sup> | 271 |
| **AaTAR3** | 1.093 × 10<sup>-7</sup> | 5.421 × 10<sup>-5</sup> | 496 |

*\*EC<sub>50</sub> values were obtained by nonlinear regression analysis of sigmoidal dose–response curves* *(Fig. 3). Relative potency (fold difference) was calculated as EC<sub>50</sub>(OA)/ EC<sub>50</sub>(TA).*

### 3.4. Co-application of biogenic amines with tyramine at AaTAR1–AaTAR3

To determine whether other biogenic amines might disrupt TA-induced receptor activation, AaTAR1–AaTAR3 were stimulated with TA (10^-5^ M) alone or in combination with OA, dopamine or serotonin (10^-5^ M each) (**Fig. 4**). Across all receptor subtypes, co-application of TA with these other biogenic amines did not significantly affect TA-evoked luminescence responses, whereas the BSA assay-medium control remained significantly lower than all ligand-stimulated responses (**Fig. 4A–C**). These findings indicate that neither OA, dopamine nor serotonin is able to detectably interfere with TA-evoked activation of any of the TA receptor subtypes (AaTAR1– AaTAR3) in *A. aegypti*.

**Figure 4.**
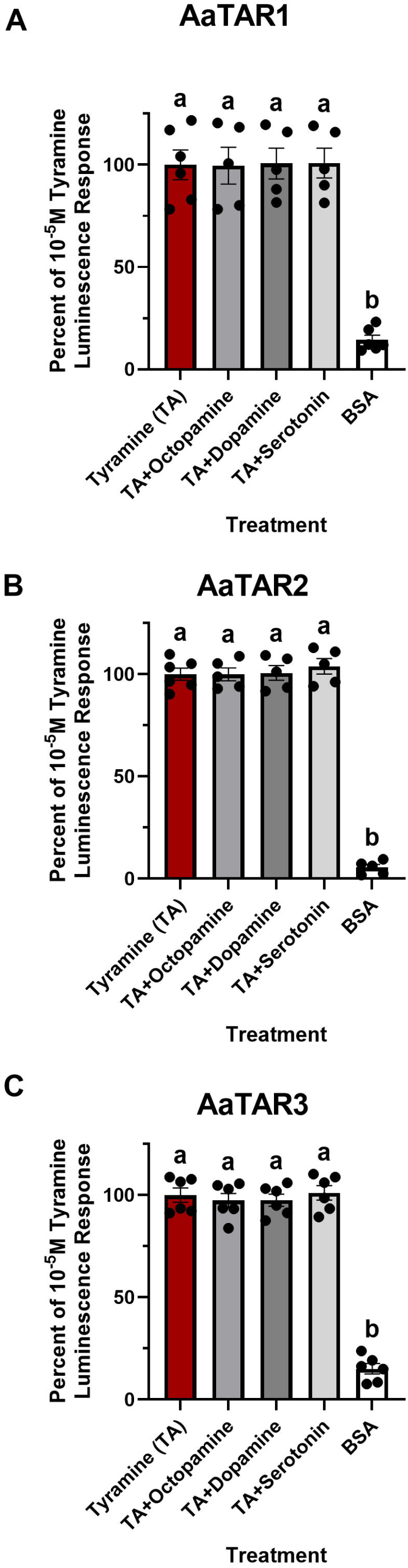
Effects of other biogenic amines on tyramine-evoked activation of AaTAR1– AaTAR3. Aequorin-CHO-K1 cells transiently expressing (**A**) AaTAR1, (**B**) AaTAR2 or (**C**) AaTAR3 were stimulated with tyramine (TA; 10^-5^ M) alone or co-applied with octopamine (OA), dopamine or serotonin (each 10^-5^ M). Luminescence responses were normalized to the TA-only condition within each receptor subtype (TA = 100%). BSA served as the negative (vehicle) control. Co-application of OA, dopamine or serotonin with TA did not significantly affect receptor activation compared to TA alone at any receptor subtype. Different letters above bars indicate statistically significant differences among treatments within each receptor subtype determined by one-way ANOVA followed by Tukey’s multiple-comparisons test; p < 0.05. Bars represent mean ± SEM (n = 5–6).

### 3.5. Antagonist Profiling of AaTAR1–AaTAR3

To evaluate subtype-specific antagonist sensitivity, AaTAR1–AaTAR3 were challenged with tyramine (TA; 10^-5^ M) alone or in the presence of individual aminergic antagonists (each 10^-5^ M; phentolamine, gramine, metoclopramide, yohimbine, mianserin or epinastine), with responses normalized to the TA-only condition (TA = 100%) (**Fig. 5**). Distinct patterns of antagonist inhibition were observed across the three AaTARs. For AaTAR1, yohimbine was the most potent antagonist, leading to significantly reduced (∼60%) TA-evoked activation, whereas gramine and mianserin produced moderate, yet significant (∼25-30%) inhibition of TA-evoked activation. Comparatively, phentolamine, metoclopramide and epinastine did not significantly change AaTAR1 responses to TA relative to controls treated with TA alone (**Fig. 5A**). Next, phentolamine produced the strongest inhibition (∼80%), while gramine also significantly reduced (∼50%) TA-evoked activation of AaTAR2. Comparatively, mianserin and epinastine caused mild, but significant inhibition (∼25%) of TA-evoked activation of AaTAR2 while metoclopramide and yohimbine did not affect AaTAR2 activation (**Fig. 5B**). AaTAR3 exhibited a comparatively narrower antagonist sensitivity profile than AaTAR1 and AaTAR2, with significant inhibition primarily observed by phentolamine (∼30%) and partial, but significant inhibition (∼20%) by gramine and epinastine. In contrast, metoclopramide, yohimbine and mianserin did not significantly alter TA-evoked activation of AaTAR3 (**Fig. 5C**). Together, these results demonstrate differential antagonist sensitivity across AaTAR1–AaTAR3, revealing pharmacological divergence among tyramine receptor subtypes in *A. aegypti*.

**Figure 5.**
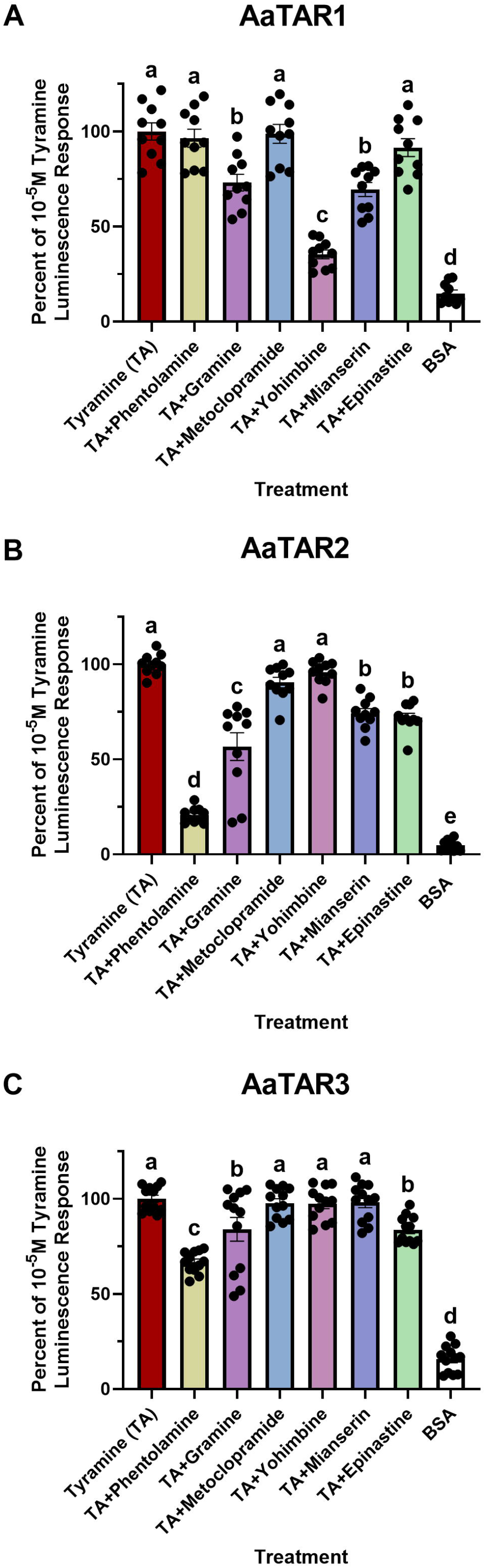
Differential antagonist sensitivity profiles of tyramine-evoked activation of AaTAR1–AaTAR3. CHO-K1 cells stably expressing aequorin and transiently expressing (A) AaTAR1, (B) AaTAR2 or (C) AaTAR3 were stimulated with tyramine (TA; 10^-5^ M) alone or co-applied with individual antagonists (10^-5^ M). Luminescence responses were normalized to the TA-only condition within each receptor subtype (TA = 100%); BSA served as the negative (vehicle) control. Different letters above bars indicate statistically significant differences among treatments within each receptor subtype determined by one-way ANOVA followed by Tukey’s multiple-comparisons test; p < 0.05. Bars represent mean ± SEM (n = 10–12).

## 4. Discussion

Tyramine receptors (TARs) are key components of insect aminergic signaling networks, modulating locomotion, metabolism, sensory processing and behaviour across a range of insect species (Roeder, 2020, 2005). Although TARs have been cloned and characterized in several insects (Blenau and Baumann, 2001; Roeder, 2005; Saudou et al., 1990), receptor subtypes can vary in ligand selectivity and pharmacological responsiveness, utilize distinct intracellular pathways and in some cases, exhibit substantial cross-reactivity with OA (Finetti et al., 2021b; Ma et al., 2019; Reale et al., 1997). However, subtype-specific functional characterization in mosquito vectors remains comparatively limited. In this study, we cloned and functionally characterized three *A. aegypti* tyramine receptor subtypes (AaTAR1–AaTAR3) and demonstrated that they share a strong agonist preference for TA, while displaying subtype-specific pharmacological divergence in antagonist sensitivity.

Agonist profiling and dose–response analyses confirmed that all three AaTAR subtypes are preferentially activated by TA over OA, dopamine and serotonin. Each receptor exhibited robust sigmoidal concentration–response relationships to TA, whereas OA displayed partial agonist activity requiring substantially higher concentrations to elicit comparatively weaker responses that were ∼50% or less of the luminescence response achieved using TA. Across the three receptor subtypes, OA was between 271–496-fold less potent than TA, underscoring pronounced ligand discrimination within the mosquito tyraminergic system. This marked potency difference reflects a high degree of functional selectivity across AaTAR subtypes. Activation of AaTAR1– AaTAR3 also produced measurable intracellular Ca^2+^ responses in the aequorin-based heterologous expression system used in this study. TARs from other insects, including *Drosophila melanogaster* and *Bombyx mori*, have been reported to couple to Gq-mediated pathways and promote IP_3_-dependent Ca^2+^ release (Huang et al., 2009; Robb et al., 1994), and a similar Ca^2+^-dependent signaling mechanism may operate in *A. aegypti*. Comparison of TA-evoked response magnitude further showed that AaTAR2 produced the strongest calcium-dependent luminescence response among the three receptor subtypes, whereas AaTAR1 and AaTAR3 displayed lower response magnitudes. The preferential activation of the three TARs by TA is broadly consistent with previous studies of insect TARs, in which TAR1 and TAR2 usually show greater selectivity for TA, with TAR2 exhibiting the highest specificity, while TAR3 has been reported to be activated by multiple biogenic amines (Bayliss et al., 2013; Blenau et al., 2017).

In *Drosophila*, TAR1 was first thought to be an octopamine receptor because it responded to both OA and TA (Arakawa et al., 1990; Saudou et al., 1990). Later, pharmacological analyses in several insects revealed TAR1 had a higher affinity for TA, leading to its reclassification as a bona fide tyramine receptor (Finetti et al., 2021a; Ohta and Ozoe, 2014). Consistent with this revised classification, AaTAR1 in *A. aegypti* exhibited a pronounced TA preference, with ∼460-fold (∼2.7 orders of magnitude) greater ligand potency in favour of TA over OA in the current concentration–response analysis. Comparable TA-preferential activation has been reported in multiple insect species, including *Rhodnius prolixus* (Hana and Lange, 2017b)*, Locusta migratoria* (Poels et al., 2001; Vanden Broeck et al., 1995) and *B. mori* (Ohta et al., 2003), where TA activates TAR1 at concentrations of roughly ∼2 orders of magnitude lower than OA. In *Chilo suppressalis*, the potency difference appears more modest, with TA reported to be ∼3-fold more potent than OA (Wu et al., 2013). In contrast, in *D. melanogaster* (Enan, 2005; Saudou et al., 1990) and *Apis mellifera* (Blenau and Baumann, 2001), the TA–OA potency gap is ∼1 order of magnitude, which nonetheless supports an overall conservation of tyramine selectivity within the TAR1 subtype.

A similar pattern of tyramine-preferential activation was observed for AaTAR2. In our dose– response analysis, TA was ∼271-fold more potent than OA in activating AaTAR2, which reinforces its classification as a functionally validated TA receptor. This level of ligand selectivity is well aligned with studies on TAR2 orthologs in other insect species, since this receptor demonstrates strong TA selectivity and activates Ca^2+^-dependent signaling pathways (Huang et al., 2009; Wu et al., 2015; Zhang and Blumenthal, 2017). Activation of TAR2 in *B. mori* and *D. melanogaster* causes intracellular Ca^2+^ mobilization with a marked preference for TA relative to OA (Bayliss et al., 2013; Cazzamali et al., 2005; Huang et al., 2009). Similar selectivity has also been reported in *C. suppressalis* whereby TA induces a substantial increase in intracellular Ca^2+^ levels, whereas OA, dopamine and serotonin fail to elicit comparable responses (Wu et al., 2015). Collectively, these results support a conserved activation profile for TAR2 as a highly selective TA receptor across insects.

Although TAR3 has been far less extensively characterized across insects compared to TAR1 and TAR2 receptor subtypes, studies in *D. melanogaster* indicate that TAR3 mediates intracellular Ca^2+^ mobilization with moderate selectivity for TA. Interestingly, *D. melanogaster* TAR3 displayed broader amine responsiveness relative to other TAR subtypes as it was activated similarly by OA and dopamine, albeit requiring concentrations ∼2-3 orders of magnitude higher relative to TA (Bayliss et al., 2013). In contrast, AaTAR3 demonstrated strong responsiveness to TA relative to OA and other tested biogenic amines, with TA eliciting ∼496-fold greater potency than OA, supporting its clear classification as a functionally selective TA receptor in *A. aegypti*.

To further investigate receptor specificity, we conducted ligand co-application experiments under saturating TA conditions (10^-5^ M) to determine whether other biogenic amines could interfere with receptor activation. Co-applying TA along with either OA, dopamine or serotonin did not change TA-evoked responses of AaTAR1–AaTAR3, with activation levels remaining comparable to treatments with TA alone. This lack of diminished receptor activation by TA in the presence of other aminergic ligands further supports a high degree of ligand selectivity across all three TA receptor subtypes in *A. aegypti*.

Subsequent antagonist profiling revealed marked divergence in pharmacological characteristics with respect to antagonist sensitivity among AaTAR1–AaTAR3. While AaTAR1 activation by TA was partially compromised by gramine and mianserin, yohimbine caused the strongest inhibitory effect on TA-induced activation and this was consistent with the conserved pharmacological profile of insect type-1 tyramine receptors (TAR1) (Finetti et al., 2021a). Comparable antagonist sensitivities have been described in *Periplaneta americana* and *Plutella xylostella*; yohimbine was the most effective antagonist, followed by mianserin (Blenau et al., 2017; Ma et al., 2019; Rotte et al., 2009). Both epinastine and phentolamine showed no antagonistic effects against AaTAR1 in line with earlier observations in *P. americana* (Blenau et al., 2017). Metoclopramide also did not cause any significant suppression TA activation of AaTAR1. Similarly, metoclopramide was only weakly active against TAR1 in *B. mori* (Ohta et al., 2003), although it produced a modest but significant inhibition of the TA-induced response of TAR1 in *R. prolixus* (Hana and Lange, 2017b).

Consistent with our results, yohimbine acts as a potent and conserved TAR1 antagonist across different insect species, including *D. melanogaster*, *L. migratoria*, *R. prolixus* and *B. mori*, which underscores the evolutionary conservation of TAR1-mediated signaling (Blenau et al., 2017; Enan, 2005; Hana and Lange, 2017b; Ohta et al., 2003; Reim et al., 2017; Robb et al., 1994; Rotte et al., 2009; Saudou et al., 1990; Vanden Broeck et al., 1995); therefore, these observations further support the pharmacological classification of AaTAR1 as a TAR1 receptor subtype. An additional important point is that there is functional evidence that extends these findings beyond the scope of the receptor pharmacology, as in adult *Laodelphax striatellus*, TAR1 antagonism via dsRNA-mediated knockdown or yohimbine injection leads to a decrease in their survival (Yan et al., 2026); this might be a potential of TARs as promising targets for novel insect control strategies.

Unlike the conserved pharmacological profile we observed for AaTAR1, AaTAR2 and AaTAR3 exhibited unique antagonist sensitivities, which may point to a possible functional variation among receptor subtypes. AaTAR2 had a clear antagonist profile, with the strongest inhibitory effect observed by phentolamine, followed by gramine, whilst AaTAR3 showed comparatively low sensitivity to the tested antagonists, indicating a narrower pharmacological profile. In *A. mellifera*, TAR2 was most potently blocked by mianserin, followed by yohimbine, but was found to be least sensitive to phentolamine (Reim et al., 2017), whereas AaTAR2 showed only partial (∼25%) antagonism by mianserin and no response to yohimbine. Strikingly, the pronounced phentolamine sensitivity observed for AaTAR2 therefore represents a unique pharmacological feature not previously reported for insect TAR2 orthologs, potentially highlighting a mosquito-specific receptor profile. In *D. melanogaster*, TAR3 has been reported to be insensitive to several classical biogenic amine antagonists, including yohimbine, mianserin, phentolamine and metoclopramide when attempting to block the effects of TA inhibition on forskolin-stimulated cAMP levels (Bayliss et al., 2013). Our AaTAR3 data also showed generally low sensitivity to the tested antagonists when measuring calcium signaling in the current study. Nonetheless, we observed partial inhibition by phentolamine that may reflect the closer evolutionary relationship betweenAaTAR2 and AaTAR3 (Finetti et al., 2023) and highlight a subtle but notable divergence in antagonist sensitivity between species. Our results reveal distinct pharmacological sensitivity profiles among the three TAR subtypes and indicate functional differences in their signaling properties.

## Conclusion

Overall, this study provides the first comprehensive functional deorphanization and pharmacological characterization of the three tyramine receptor subtypes (AaTAR1–AaTAR3) in *A. aegypti*. All three of these receptors revealed strong and highly selective activation by TA, with minimal or no responses to other biogenic amines. This confirms their high specificity and establishes TA as their endogenous aminergic ligand. Antagonist profiling determined clear subtype-specific sensitivity patterns, demonstrating that AaTAR1–AaTAR3 are pharmacologically distinct rather than functionally redundant. Taken together, these findings validate AaTAR1–AaTAR3 as bona fide TA receptors and provide new insights into the organization of mosquito tyraminergic signaling, pointing to the functional diversity of TAR subtypes. Importantly, these subtype specific pharmacological profiles could position *A. aegypti* TA receptors as promising targets for the development of selective next generation vector control strategies. Future studies integrating in vivo functional analyses, immunohistochemical localization and structural modeling will be essential to elucidate receptor specific functions and enable the rational design of targeted modulators of mosquito tyraminergic signaling.

## Footnotes

### Author Contributions

Conceptualization: S.A., J.-P.P.; Methodology: S.A., J.-P.P.; Validation: S.A., J.-P.P.; Formal analysis: S.A., J.-P.P.; Investigation: S.A., J.-P.P.; Resources: J.-P.P.; Data curation: S.A., J.-P.P.; Writing - original draft: S.A.; Writing - review & editing: S.A., J.-P.P.; Visualization: S.A., J.-P.P.; Supervision: J.-P.P.; Project administration: J.-P.P.; Funding acquisition: J.-P.P.

### Data and Resource Availability

All data supporting the findings of this study are available within the article.

